## Supplements for "Beyond fear centers - a distributed fMRI-based neuromarker for the subjective experience of fear"

**Zhou et al.**

### Supplementary Methods

#### MRI data acquisition

Functional MRI data for the discovery and validation paradigms was acquired using a T2*-weighted echo-planar imaging (EPI) pulse sequence (repetition time = 2s, echo time = 30ms, 36 slices, slice thickness = 3.8mm, no gap, field of view = 200 × 200mm, resolution = 64 × 64, flip angle = 90°, voxel size = 3.125 × 3.125 × 3.8mm). To improve spatial normalization and exclude participants with apparent brain pathologies a high-resolution T1-weighted image was acquired using a 3D spoiled gradient recalled (SPGR) sequence (repetition time = 8ms, echo time = 3ms, 176 slices, slice thickness = 1mm, no gap, field of view = 256 × 256mm, acquisition matrix = 256 × 256, flip angle = 8°, voxel size = 1 × 1 × 1mm).

#### Threat conditioning paradigms

In Reddan, et al. [1] study, 68 healthy subjects (45 females, mean ± SD age = 29.64 ± 15.89 years) underwent an auditory threat-conditioning paradigm inside of the fMRI environment. The CSs were 2 pure tones (800 and 170HZ) and the US (unconditioned stimulus) was a mild electric shock (200ms duration, 50 pulses/s) to the right wrist. One of the tones (counterbalanced across participants) was served as the CS+ and was paired with a shock on 33% of the trials while the other tone was designated as CS- and was never paired with a shock. Acquisition consisted of 8 CS+ tones paired with shock, 8 unreinforced CS+ tones, and 8 CS- tones. Each CS was presented for 4s and the interstimulus interval was 10s. For paradigm, MRI data acquisition and preprocessing details please see Reddan, et al. [1].

The visual threat conditioning dataset included 58 male subjects (mean ± SD age = 20.68 ± 1.76 years) who exhibited average unreinforced CS+ SCR > CS- SCR (skin conductance response) during acquisition underwent a visual threat-conditioning paradigm during fMRI scanning [2]. Briefly, one colored square (CS+, 4s) coincided with a mild electric shock (US, 2ms) to the right wrist with 43% contingency, whereas the other differentially colored square (CS-, 4s) was never paired with the US. Acquisition included two runs and each run contained 8 non-reinforced presentations of the CS+ and the CS-, intermixed with an additional 6 presentations of the CS+ paired with the shock (CS+U). Stimuli were presented in a pseudorandom order with a 9-12s interstimulus interval. Two color sets were used and balanced across subjects to control for effects of the designated colors (color set A: red = CS+, blue = CS-; color set B: Green = CS+, pink = CS-). For paradigm, MRI data acquisition details please see Zhou et al., 2019. Of note, in this study we re-preprocessed the fMRI data using the same pipeline that the fear datasets used.

For both datasets the subject-level GLM analyses included 3 boxcar regressors: reinforced CS+, unreinforced CS+ and CS- and in this study we only analyzed beta estimates for unreinforced CS+ and CS- conditions.

### Supplementary results

#### VIFS requires, but not fully depends on, the visual cortex

To test how strongly the VIFS' performance depended on visual cortex, which might encode emotion schemas or aspects of high-level visual processing related to emotional experience [3,4], we re-trained the fear decoder excluding the occipital lobe. We found that the overall prediction-outcome correlations (discovery, cross-validated r = 0.51, 95% CI = [0.43, 0.58]; validation, r = 0.57, 95% CI = [0.45, 0.66]) and classification accuracies were slightly decreased but were still comparable (see **Supplementary Fig. 2** for details), suggesting that the fear-predictive signals might be partly embedded in the visual cortex but the contribution of visual cortical patterns is small.

**VIFS outperformed two other related patterns**

Previous studies have developed two related brain signatures (V.T. decoder and PINES) to predict subjective fear ratings [5] and general negative emotion experience [3], respectively. We compare the performance of the VIFS with the V.T. decoder on the discovery, validation and generalization cohorts. Note that (1) when applying the VIFS to the discovery cohort and the V.T. decoder to the generalization cohort we used cross-validation procedures to get unbiased estimates, (2) we retrained the V.T. model using labels 1-6 instead of 0-5 for display purposes (of note, this procedure changes only the intercept/bias but not the pattern weights of the predictive model and has no effects on the prediction-outcome correlation or the forced-choice classification) and (3) given that within-subject mean centering could inflate the overall prediction-outcome correlation we did not demean the samples when developing and testing V.T. decoder on the generalization cohort (for the other comparisons we used the V.T. decoder developed on the within-subject mean centered samples as the authors did in their study). We found that although statistically significant, the prediction-outcome correlation coefficients (r = 0.18, 95% CI = [0.07, 0.29], permutation test one-tailed P = 0.011; r = 0.26, 95% CI = [0.07, 0.42], permutation test one-tailed P = 0.004) of the V.T. decoder on the discovery and validation cohorts were very low (Fig 3C) as compared with VIFS' predictions (r = 0.57, 95% CI = [0.49, 0.63]and r = 0.59, 95% CI = [0.45, 0.65], respectively) (Fig 3D); on the other hand, the VIFS predicted the generalization cohort comparably to the V.T. decoder (r = 0.56, 95% CI = [0.49, 0.63], permutation test one-tailed P < 0.001; r = 0.64, 95% CI = [0.54, 0.71], respectively). We additionally trained the V.T. decoder without mean-centering and found that the prediction correlations on the discovery and validation cohorts were similar to the centered model. Moreover, the V.T. decoder failed to classify high versus moderate subjective fear in the discovery and validation cohorts as well as moderate versus low subjective fear in the validation cohort while the VIFS accurately classified low, moderate and high subjective fear in all three cohorts (all accuracies > 80%. Table 1).

Furthermore, we found that the VIFS predicted subjective fear considerably better than the PINES on all of the three fear cohorts while the PINES was more sensitive to predicted general negative emotion experience as compared with the VIFS (see Table 2 for details). In support of the prediction-outcome correlations, VIFS predicted high versus low subjective fear more accurately than the PINES in all of the three fear cohorts while PINES could better distinguish high versus low negative emotions in the PINES holdout dataset ($n = 61$) in terms of forced-choice classification accuracy and effect size (Fig. 6A). Importantly, we confirmed that only 2 stimuli from IAPS (International Affective Picture System) were used in both studies, arguing against that the significant predictions were due to the overlapping stimuli.

**VIFS responses mediate fear induced by negative emotion**

We employed multilevel mediation analyses to investigate the associations among VIFS response, PINES response and fear rating. Firstly, we tested whether the VIFS response could explain the association between the PINES response and subjective fear rating. We found that in the discovery cohort the PINES response was positively correlated with the VIFS response (path a; $\beta = 0.55$, 95% CI = [0.47, 0.64]; $Z = 4.21$, $P < 0.001$; Cohen's $d = 0.38$, the VIFS response had a positive association with fear rating independently of the PINES response (path b; $\beta = 0.19$, 95% CI = [0.18, 0.21]; $Z = 4.49$, $P < 0.001$; Cohen's $d = 0.37$) and there was a significant mediation effect (path a×b; $\beta = 0.09$, 95% CI = [0.08, 0.11]; $Z = 4.13$, $P < 0.001$; Cohen's $d = 0.21$). Together with the significant path c ($\beta = 0.15$, 95% CI = [0.12, 0.18]; $Z = 4.43$, $P < 0.001$; Cohen's $d = 0.18$) and path c' ($\beta = 0.04$, 95% CI = [0.01, 0.07]; $Z = 2.99$, $P = 0.003$; Cohen's $d = 0.05$) our results suggested that the VIFS response partially mediated the effect of PINES response on the fear rating (Fig. 6B). These findings were replicated in the validation cohort.

In addition, we conducted another multilevel mediation analysis with VIFS response as x, PINES response as m and fear rating as y. We found that the PINES could not mediated VIFS response – fear rating association in the discovery cohort (Fig. 6C) and in the validation cohort although the mediation effect was significant the mediation effect size was much smaller as compared with the above model (Cohen's d = 0.06 versus Cohen's d = 0.21). Thus, the PINES response was unlikely to mediate the VIFS response effect on the fear rating.

**Post hoc exploration of predictive models**

To test whether the hyperparameter optimization (i.e., the "C" parameter) could improve the VIFS performance we performed additional post hoc analyses optimizing the C parameter (across C = [0.0001, 0.001, 0.01, 0.1, 1, 10]) using nested cross-validation. This procedure did not lead to strong improvements in the discovery cohort (C = 1 prediction-outcome correlation r = 0.5647; cross-validated C r = 0.5648). We additionally chose the optimal C parameter using the whole discovery cohort (evaluated by repeated 10-fold cross validation) and predicted the validation and generalization cohorts with the best model. We found that it predicted the validation cohort slightly better (r = 0.5995 vs. 0.5926 with C = 1), but predicted the generalization cohort slightly worse (r = 0.5566 vs. 0.5627 with C = 1). In line with previous studies, our findings suggest that the performances of the SVR models are often insensitive to the choice of C as long as C is in an intermediate range.

In addition, we performed a recursive feature elimination (RFE) analysis in which the importance of a feature was estimated by the absolute value of its corresponding predictive weight, and less important features (i.e., with low predictive weights) were eliminated recursively. We removed 10,000 voxels each time and re-trained the model with the remaining voxels and predicted validation and generalization cohorts until less than 10,000 voxels were left after the elimination (we further trained the model with the most important 10,000 voxels). We found that as the number of voxels/features decreased the prediction-outcome correlation of the validation cohort also decreased, but that predictive accuracy for the generalization cohort slightly increased until around 200,000 out of 225,672 voxels were removed (Supplementary Fig. 5A). Moreover, Moreover, in line with a previous study [6] we plotted the final predictive SVR features after the RFE procedure, with the final number of features = 20,000 (Supplementary Fig. 5B). The most compact set of informative voxels was very similar to the reliable voxels revealed by the bootstrap test, suggesting that the informative voxels were consistent across analytical approaches.

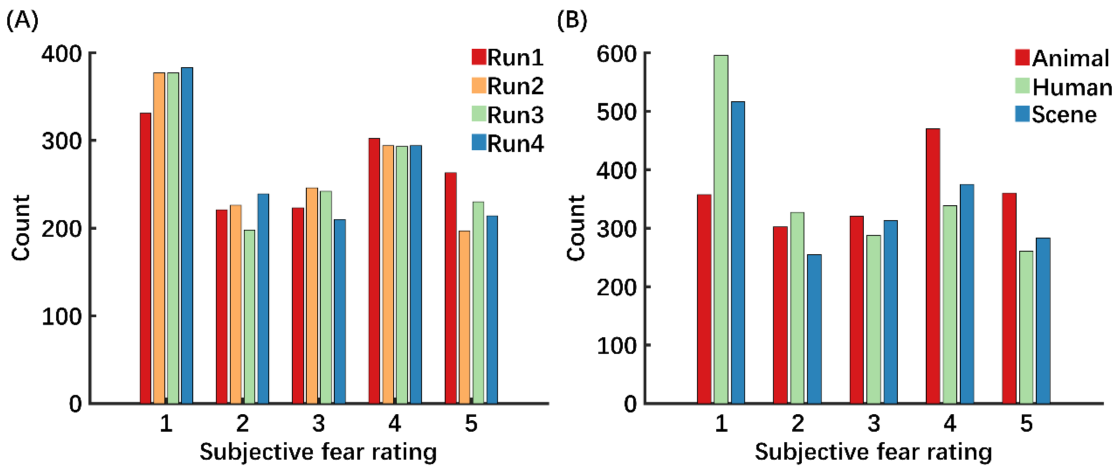


**Supplementary Fig. 1 Self-reported fear levels are generally evenly distributed across categories and runs.** Panels A and B depict the distribution of the subjective fear ratings in each run and for each semantic content (i.e., animal, human and scene), respectively.

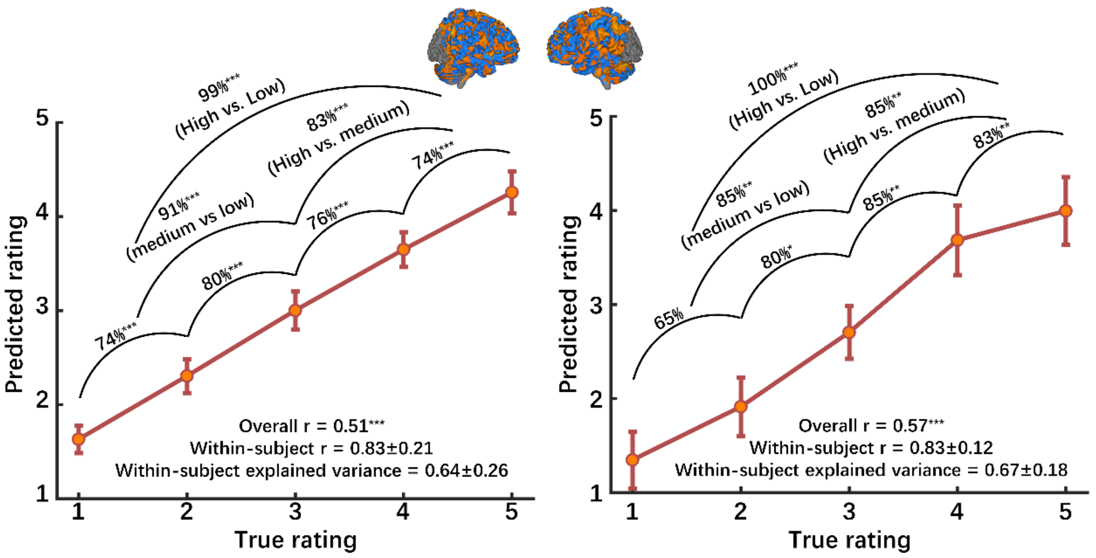


**Supplementary Fig. 2 VIFS response pattern response without occipital lobe.**

Panels A and B depict the predicted fear experience (subjective ratings) compared to the actual level of fear for the cross-validated discovery cohort and the independent validation cohort, respectively. Accuracies reflect forced-choice comparisons.

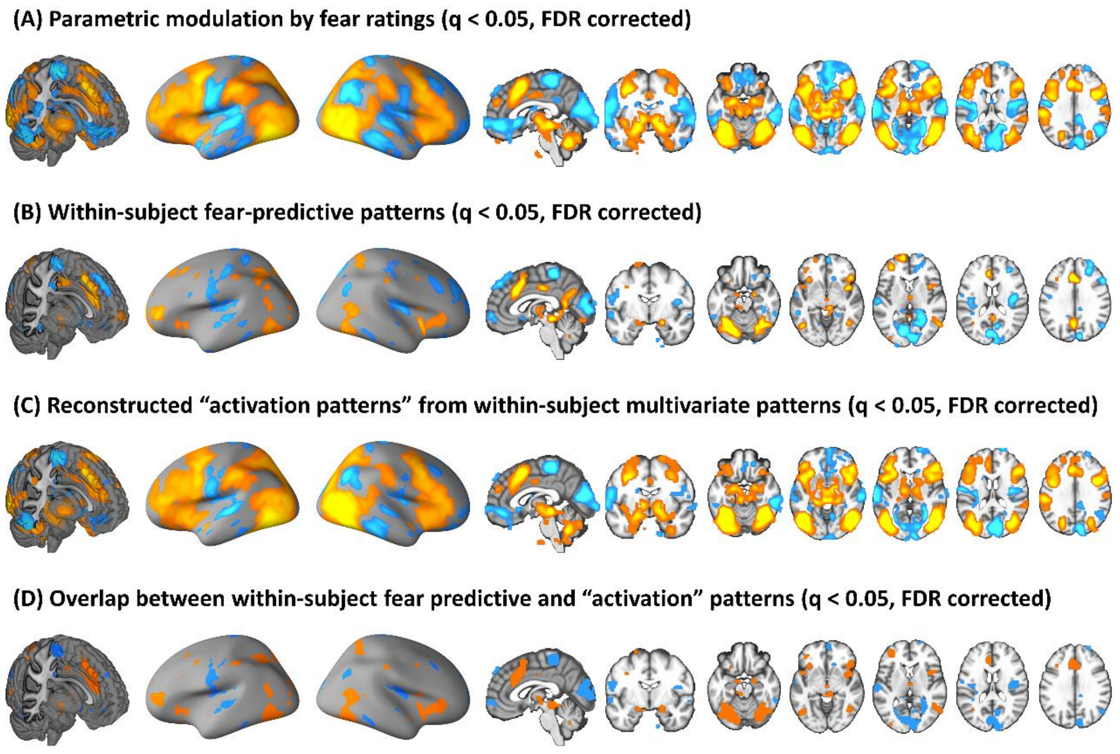


**Supplementary Fig. 3 Subjective experience of fear is associated with and predicted by distributed brain regions.** Panel A shows univariate parametric effects of fear ratings. Panel B summaries multivariate patterns trained on individual subjects and depicts brain regions consistently predictive of subjective fear across participants. Panel C shows thresholded transformed ‘activation patterns’ from within-subject fear-predictive patterns. Panel D demonstrates overlapping (i.e., from a conjunction analysis) brain regions between panel B and C. Hot color indicates positive associations (panels A, C) or weights (panels B) where as cold color indicates negative associations (panels A, C) or weights (panels B).

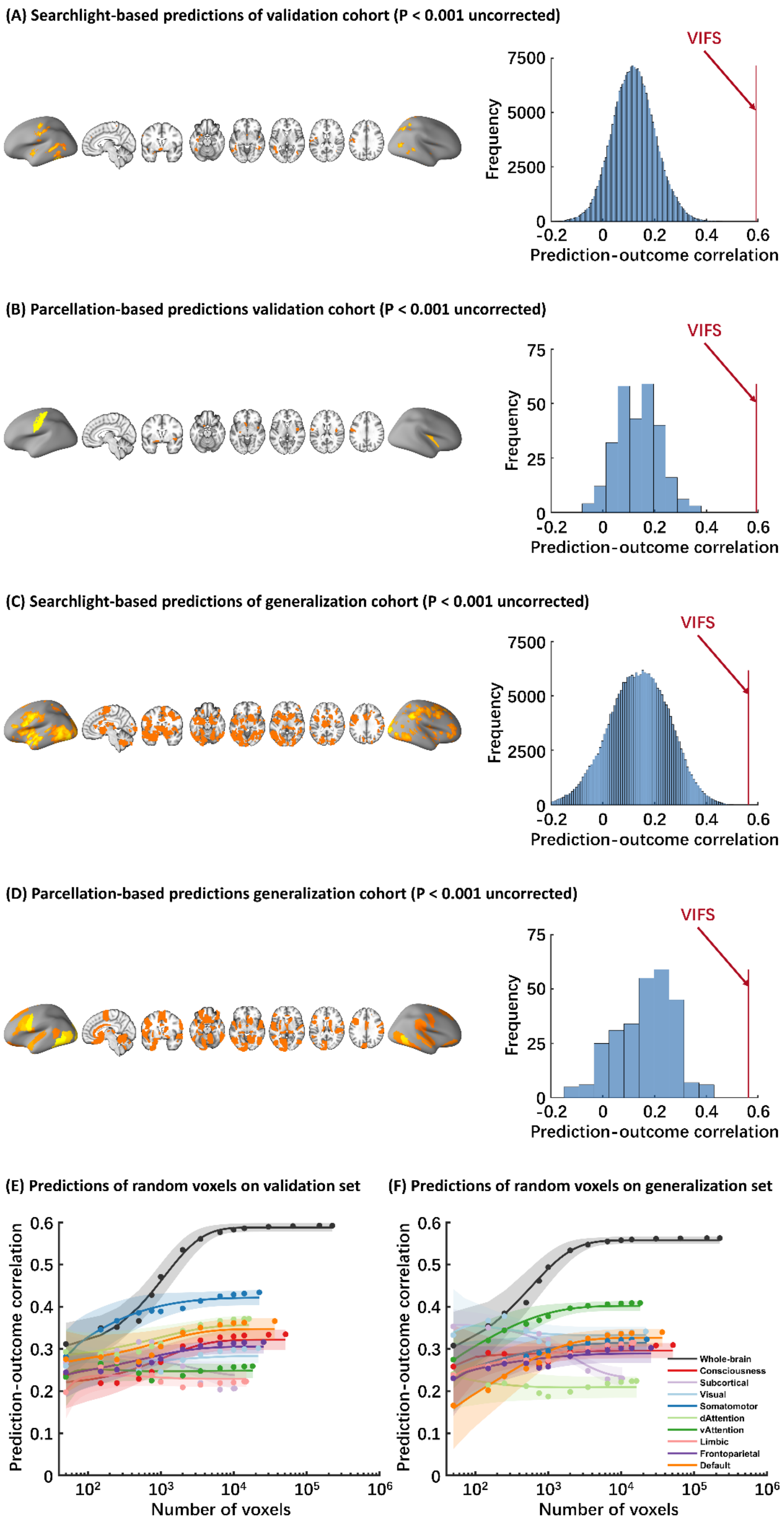

(A) Searchlight-based predictions of validation cohort (P < 0.001 uncorrected)
VIFS
Frequency
Prediction-outcome correlation
(B) Parcellation-based predictions validation cohort (P < 0.001 uncorrected)
VIFS
Frequency
Prediction-outcome correlation
(C) Searchlight-based predictions of generalization cohort (P < 0.001 uncorrected)
VIFS
Frequency
Prediction-outcome correlation
(D) Parcellation-based predictions generalization cohort (P < 0.001 uncorrected)
VIFS
Frequency
Prediction-outcome correlation
(E) Predictions of random voxels on validation set
Prediction-outcome correlation
Number of voxels
(F) Predictions of random voxels on generalization set
Prediction-outcome correlation
Number of voxels
Whole-brain
Consciousness
Subcortical
Visual
Somatomotor
dAttention
vAttention
Limbic
Frontoparietal
Default

**Supplementary Fig. 4 Predictions of models trained on discovery cohort on validation and generalization cohorts.** Panel A and B show brain regions that can significantly predict subjective fear ratings of the validation cohort revealed by searchlight- and parcellation-based analyses. Panel C and D show brain regions that can significantly predict subjective fear ratings of the generalization cohort revealed by searchlight- and parcellation-based analyses. Histograms: cross-validated predictions (correlations) from searchlights or parcellations. Red line indicates the prediction-outcome correlation from VIFS. In line with the cross-validated predictions on the discovery cohort all images are thresholded at $P < 0.001$ uncorrected for display purpose. Panel E and F demonstrate that the information about subjective experience of fear is distributed across multiple systems. Model performance was evaluated as increasing numbers of voxels/features (x axis) were used to predict subjective fear in different regions of interest including the entire brain (black), consciousness network (red), subcortical regions (light purple) or large-scale resting-state networks. The models were always trained with the discovery dataset and the prediction-outcome correlations in the (E) validation and (F) generalization cohorts were evaluated with the same selected voxels. Colored dots indicate the correlation coefficients, solid lines indicate the mean parametric fit and shaded regions indicate standard deviation. Model performance is optimized when approximately 10,000 voxels are randomly sampled across the whole-brain.

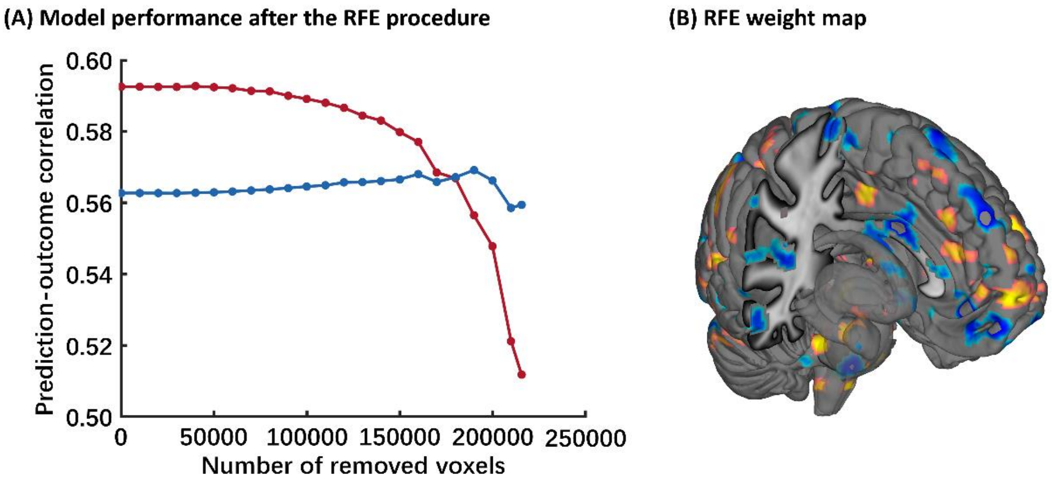


**Supplementary Fig. 5 Model performance and SVR features after the recursive feature elimination procedure.** (A) Model performance after each elimination (10,000 voxels). Red line indicates the predictions on validation cohort and the blue line indicates the predictions on generalization cohort. (B) Weight map showing the final predictive SVR features after the RFE procedure, with the final number of features = 20,000.

Supplementary Table 1 Prediction-outcome correlations (mean and std) with various numbers of voxels

| Number of voxels | Vis | SM | dA | vA | Limb | FP | DMN | Cons | subC | Whole-brain |
|---|---|---|---|---|---|---|---|---|---|---|
| 50 | 0.260 (0.037) | 0.303 (0.043) | 0.293 (0.04) | 0.267 (0.042) | 0.191 (0.046) | 0.305 (0.041) | 0.270 (0.047) | 0.295 (0.043) | 0.270 (0.046) | 0.312 (0.049) |
| 150 | 0.260 (0.033) | 0.335 (0.04) | 0.305 (0.037) | 0.300 (0.043) | 0.215 (0.044) | 0.313 (0.04) | 0.285 (0.041) | 0.304 (0.042) | 0.317 (0.039) | 0.348 (0.045) |
| 250 | 0.244 (0.028) | 0.333 (0.038) | 0.303 (0.033) | 0.311 (0.038) | 0.223 (0.038) | 0.304 (0.036) | 0.290 (0.039) | 0.301 (0.039) | 0.324 (0.030) | 0.349 (0.042) |
| 500 | 0.217 (0.021) | 0.316 (0.031) | 0.299 (0.029) | 0.302 (0.031) | 0.226 (0.032) | 0.295 (0.031) | 0.295 (0.033) | 0.294 (0.033) | 0.304 (0.022) | 0.366 (0.041) |
| 750 | 0.213* (0.019) | 0.309 (0.028) | 0.289 (0.025) | 0.294 (0.026) | 0.230* (0.029) | 0.289 (0.027) | 0.309 (0.031) | 0.302 (0.029) | 0.278* (0.019) | 0.427 (0.037) |
| 1000 | 0.216* (0.016) | 0.308* (0.025) | 0.277* (0.021) | 0.296* (0.023) | 0.233* (0.025) | 0.296* (0.024) | 0.327 (0.027) | 0.321 (0.026) | 0.259* (0.017) | 0.462 (0.033) |
| 2000 | 0.232* (0.011) | 0.323* (0.018) | 0.284* (0.016) | 0.315* (0.015) | 0.247* (0.02) | 0.316* (0.017) | 0.353* (0.018) | 0.351* (0.019) | 0.225* (0.013) | 0.513 (0.025) |
| 3500 | 0.239* (0.008) | 0.334* (0.013) | 0.292* (0.012) | 0.323* (0.011) | 0.257* (0.015) | 0.324* (0.012) | 0.362* (0.013) | 0.362* (0.014) | 0.209* (0.009) | 0.537 (0.019) |
| 6500 | 0.243* (0.006) | 0.340* (0.009) | 0.297* (0.007) | 0.328 (0.007) | 0.262* (0.009) | 0.328 (0.008) | 0.368 (0.009) | 0.368* (0.01) | 0.212* (0.005) | 0.550 (0.014) |
| 10000 | 0.244* (0.004) | 0.342* (0.006) | 0.299* (0.004) | 0.329* (0.005) | 0.264* (0.006) | 0.330* (0.006) | 0.371* (0.007) | 0.370* (0.008) | 0.214* (0.001) | 0.557 (0.011) |
| 14000 | 0.245* (0.003) | 0.343* (0.004) | 0.300* (0.002) | 0.330* (0.003) | 0.265* (0.002) | 0.331* (0.004) | 0.372* (0.006) | 0.372* (0.006) | NA | 0.559 (0.009) |
| 30000 | NA | NA | NA | NA | NA | NA | NA | 0.373* (0.003) | NA | 0.564 (0.006) |
| 65000 | NA | NA | NA | NA | NA | NA | NA | NA | NA | 0.566 (0.004) |
| 150000 | NA | NA | NA | NA | NA | NA | NA | NA | NA | 0.567 (0.002) |
| Full | 0.246 | 0.344 | 0.300 | 0.331 | 0.265 | 0.331 | 0.373 | 0.374 | 0.214 | 0.567 |

Vis, visual network; SM, somatomotor network; dA, dorsal attention network; vA, ventral attention network; Limb, limbic network; FP, frontoparietal network; DMN, default model network; Cons, consciousness network; subC, subcortical network; NA, not applicable. *indicates that the prediction is significant lower as compared with the whole-brain model using the same number of voxels (two-tailed Z-test; $P<0.05$, Bonferroni corrected).
